# Exploring mutational possibilities of KPC variants to reach high level resistance to cefiderocol

**DOI:** 10.1101/2025.02.14.638246

**Authors:** Sidonie Hanna, Kevin La, Yutaka Yoshii, Maud Gits-Muselli, Imane El Meouche, Yasmine Benhadid-brahmi, Stéphane Bonacorsi, André Birgy

**Author notes:** Corresponding Author: André Birgy.

## Abstract

**Introduction:** *Klebsiella pneumoniae carbapenemase (KPC)* is one of the most widespread carbapenemases, with over 242 variants identified. Some, like KPC-33 (D179Y), resist both ceftazidime-avibactam (CZA) and cefiderocol but show only a moderate cefiderocol MIC increase. KPC’s success stems from its genetic adaptability, enabling mutation accumulation and necessitating proactive resistance monitoring. This study investigates the mutational possibilities for three KPC variants to develop high-level cefiderocol resistance.

**Method:** Using random mutagenesis and enrichment selection over a 10-day cefiderocol exposure at increasing concentration, we investigated the mutational potential of KPC-2, KPC-3, and KPC-33 to develop high-level cefiderocol resistance. Resistance mechanisms were then analyzed through phenotypic testing and sequencing.

**Results:** Random mutagenesis generated 10⁵, 10⁴, and 10^5^ mutants for *bla*_KPC-2_, *bla*_KPC-3_, and *bla*_KPC-33_, respectively. After ten days of increasing cefiderocol exposure, MICs rose significantly, with KPC-2 mutants reaching >32 mg/L. Phenotypic analysis revealed a common resistance profile across all tested mutants, with resistance to ceftazidime, ceftazidime-avibactam, cefixime, and piperacillin, but restored susceptibility to carbapenems and most other β-lactams. Sequencing identified key mutations (D179Y- D209V in KPC-2, L169P in KPC-3), while chromosomal changes, particularly *cir*A and *ybi*X disruptions, played a crucial role in resistance evolution.

**Discussion:** Our results show that the enriched mutations in KPC genes are not sufficient to confer high cefiderocol MICs, suggesting that, in the studied variants, no simple mutational pathway allows the KPC enzyme to efficiently hydrolyze cefiderocol. These findings underscore the interplay between enzymatic and iron transport mutations in cefiderocol resistance, highlighting the importance of surveillance to anticipate emerging resistance in KPC-producing pathogens.

---

*Klebsiella pneumoniae* carbapenemase (KPC) stands as the most prevalent carbapenemase enzyme globally, with an endemic presence across North and South America, China, Israel, Greece, and Italy (1, 2). KPC exhibits a wide-ranging resistance profile, encompassing most beta-lactams including carbapenems and classical beta-lactamase inhibitors. New beta-lactamase inhibitors are effective against KPC beta-lactamase (avibactam, vaborbactam, relebactam …) and showed effectiveness against KPC-producing Enterobacterales(3–5). However, KPC diversified rapidly with 242 clinical variants described to date (January 2025). The clinical success of KPC variants is attributed to intricate molecular mechanisms and a remarkable genetic adaptability, though our understanding of these processes remains incomplete (6, 7). While the combination of ceftazidime-avibactam (CZA) is recommended as the first-line treatment for systemic infections caused by KPC-producing Enterobacteriaceae (KPE), many KPC variants have been reported to be resistant to CZA (6, 8, 9). Recent studies highlight the emergence of clinical variants that confer cross-resistance to newly introduced therapeutic agents such as meropenem-vaborbactam (MEV) or imipenem-relebactam (IMR) (10–12). In this context, Cefiderocol has become a therapeutic option since its introduction in 2020 (9).

Cefiderocol is a siderophore cephalosporin sharing side chains similar to those of both cefepime and ceftazidime conferring stability against most beta-lactamases (13–15). The chlorocatechol group binds covalently to extracellular Fe3+, allowing its fast and active transport into the bacterial periplasm by iron TonB-dependent transporters (TBDT) such as FepA, Fiu or CirA (13, 14, 16). However, recently cross-resistance to both CZA and Cefiderocol has been described in some KPC variants and notably in KPC-33 (D179Y) and KPC-31 (D179Y-H274) (17–21). The D179Y mutation is observed in 26 out of 242 (11%) of KPC variants with KPC-33 and KPC-31, being prominent in global epidemiology (6). The structural analogy between Cefiderocol and Ceftazidime could be one of the origins of this cross-resistance, corroborated by the identification of common molecular and enzymatic mechanisms (18, 22). While described mutants currently confer a modest increase in cefiderocol MICs – close to EUCAST breackpoints (2 mg/L), it is possible that the natural evolution of KPC-2 or one of it’s variant, along with the accumulation of mutational events with time, could increase the level of resistance in the future.

This evolutionary adaptability, similar to the one observed with TEM beta-lactamases and 3^rd^ generation cephalosporins would pose an escalating threat to healthcare systems, underscoring the need to anticipate future resistance (23). In this context, we aimed to test the evolutionary possibilities for 3 KPC variants (KPC-2 and KPC-3 being the most frequent and KPC-33: a variant with increased cefiderocol MIC) to confer a high level of resistance to cefiderocol. To achieve this, we generated KPC mutant libraries through random mutagenesis to assess the potential for selecting a combination of mutations that confer high levels of resistance to Cefiderocol.

## Results

### Mutant libraries from random mutagenesis experiment

Libraries of mutants were generated by random mutagenesis experiment, resulting in a total of 1.3 x 10⁵ colonies from *bla*_KPC*-*2_, 3 x 10⁴ colonies from *bla*_KPC-3_, and 10^5^ colonies from *bla*_KPC-33_. Colonies were scraped off the agar into LB-glycerol (40%), and this cell suspension was aliquoted and stored at −80°C after thorough mixing. The average mutagenesis efficiency resulted in 3.6 substitution-type mutations per allele. It was determined on 100 randomly chosen colonies before selection on which the KPC gene was sanger sequenced.

### *In vitro* enrichment experiment and selection of mutants

Cefiderocol MICs of the ancestral variants *bla*_KPC-2,_ *bla*_KPC-3_ and *bla*_KPC-33_ were 0.25, 0.12, and 2 mg/L respectively. The increase of MICs over the ten-day exposure period is shown in Figure 1. By day 10, an increase in MICs was observed for all three exposed mutant libraries: >32, 2 and 8 mg/L corresponding to an increase by >128-fold, 17-fold, and 4-fold respectively for the libraries derived from *bla*_KPC-2_, *bla*_KPC-3_ and *bla*_KPC-33_. Starting from the well that grew at the highest cefiderocol concentration, we re-isolated 10 µL of culture by plating on LB-tetracycline. Ten colonies from each of these enrichment experiments were randomly selected for phenotypic characterization.

**Figure 1.**
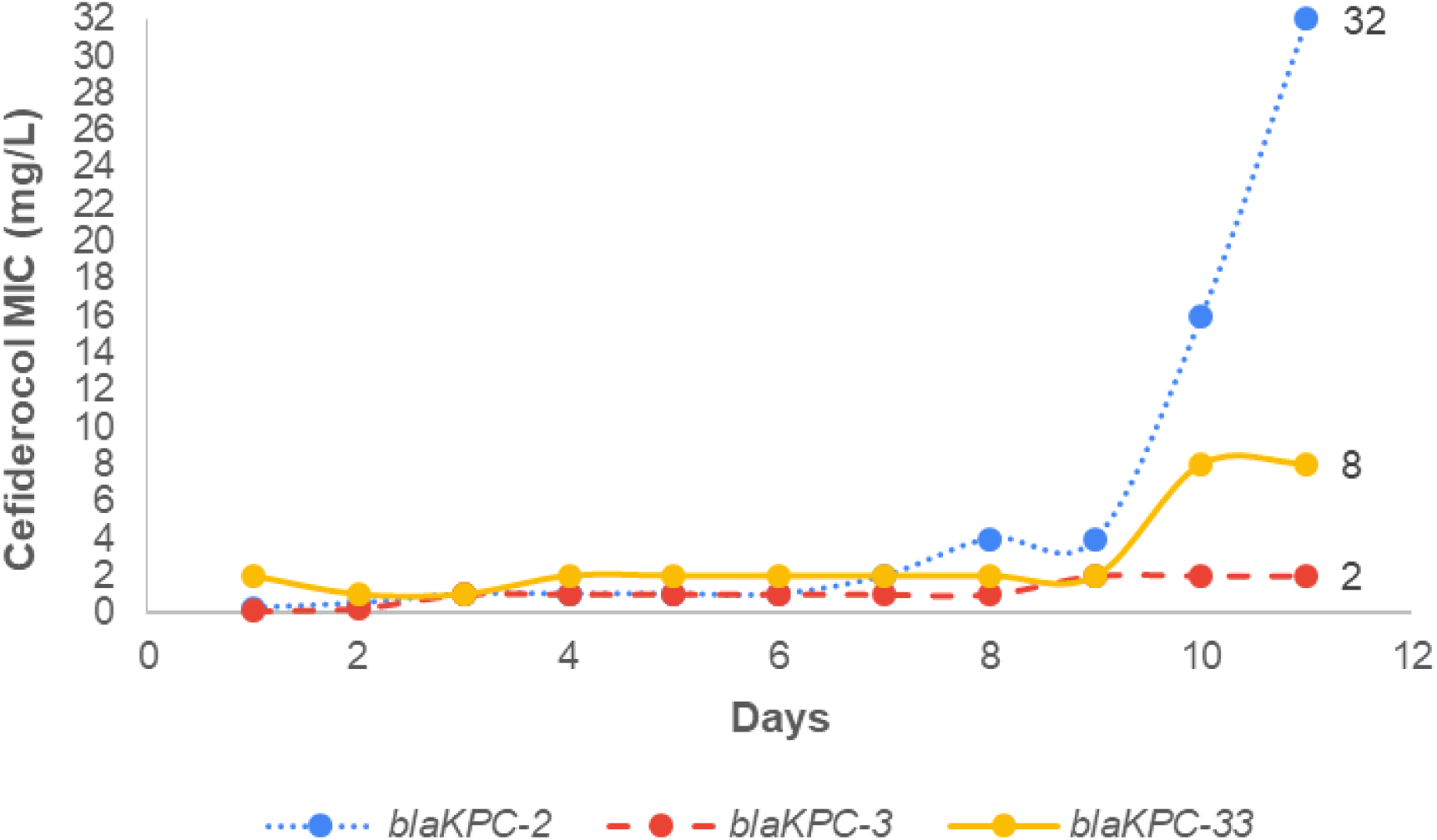
Results of the enrichment experiment of mutant libraries under increasing concentrations of cefiderocol (10 days of evolution)

Phenotypic characterization: Antibiotic susceptibility testing allowed the identification of a common resistance phenotypic profile for all tested mutants after the ten-day exposure to increasing concentrations of cefiderocol. Mutants were all resistant to ceftazidime and CZA, cefixime, and piperacillin but were susceptible to carbapenems (Table 1, Supplementary table 2 and Figure 2). So, mutations enriched in our experiment (especially from KPC-2 and KPC-3) modified the resistance capacity of the KPC protein. Also, the Cefiderocol MICs increased differently depending on the initial variant (table 1).

**Table 1.** MIC determination for selected mutants against different beta-lactams. AMP, Ampicillin ; CEP, Cephalothin, FAZ, Ceftazolin ; FOX, Cefoxitin ; AXO, Ceftriaxone ; CTX, Cefotaxime ; POD, Cefpodoxime ; FEP, Cefepime ; CAZ, Ceftazidime ; CZA; Ceftazidime-avibactam ; T/C, Ceftazdime/clavulanate ; MEM, Meropenem ; IMP, Imipenem ; PTZ, Piperacillin/tazobactam ; F/C, Cefotaxime/clavulanate; FDC, Cefiderocol

|  |  |  |  | MIC (mg/L) |  |  |  |  |  |  |  |  |  |  |  |  |  |  |  |
| --- | --- | --- | --- | --- | --- | --- | --- | --- | --- | --- | --- | --- | --- | --- | --- | --- | --- | --- | --- |
| Mutants | Ancestral allele | Ancestral mutation | Acquired mutations | AMP | CEP | FAZ | FOX | AXO | CTX | POD | FEP | CAZ | CZA | T/C | MEM | IMP | PTZ | F/C | FDC |
| KPC-2 | - | - |  | >16 | >16 | >16 | 16 | 32 | 16 | >32 | 4 | 16 | 0.5 | 16 | 4 | 2 | >64 | 2 | **0.25** |
| KPC-3 | - | H274Y |  | >16 | >16 | >16 | 16 | 32 | 16 | >32 | 8 | 64 | 1 | 32 | 4 | 2 | >64 | 4 | **0.12** |
| KPC-33 | - | D179Y |  | <8 | 16 | <8 | 8 | 8 | 4 | 8 | <1 | 128 | 24 | 2 | <1 | <0.5 | <4 | <0.12 | **2** |
| KPC-2n2 | KPC-2 | - | D179Y D209V | <8 | >16 | <8 | 8 | 8 | 4 | 16 | <1 | 128 | 32 | 2 | <1 | <0.5 | <4 | 0.5 | **>32** |
| KPC-3n4 | KPC-3 | H274Y | L169P | 16 | 16 | <8 | 8 | 8 | 4 | 16 | 2 | >128 | 12 | 2 | <1 | <0.5 | <4 | <0.12 | **2** |
| KPC-33n6 | KPC-33 | D179Y | R6H F20L | <8 | 16 | <8 | 8 | 8 | 8 | 16 | <1 | >128 | 64 | 2 | <1 | <0.5 | <4 | <0.12 | **8** |
| plasmid KPC-2n2 in fresh *E. coli* Top10 | KPC-2 | - | D179Y D209V | <8 | >16 | <8 | 8 | 8 | 4 | 16 | <1 | 128 | 32 | 2 | <1 | <0.5 | <4 | 0.25 | **2** |
| plasmid KPC-3n4 in fresh *E. coli* Top10 | KPC-3 | H274Y | L169P | 16 | 16 | <8 | 8 | 8 | 4 | 16 | 2 | >128 | 12 | 2 | <1 | <0.5 | <4 | <0.12 | **0.5** |
| plasmid KPC-33n6 in fresh *E. coli* Top10 | KPC-33 | D179Y | R6H F20L | <8 | 16 | <8 | 8 | 8 | 8 | 16 | <1 | >128 | 64 | 2 | <1 | <0.5 | <4 | <0.12 | **2** |

**Figure 2:**
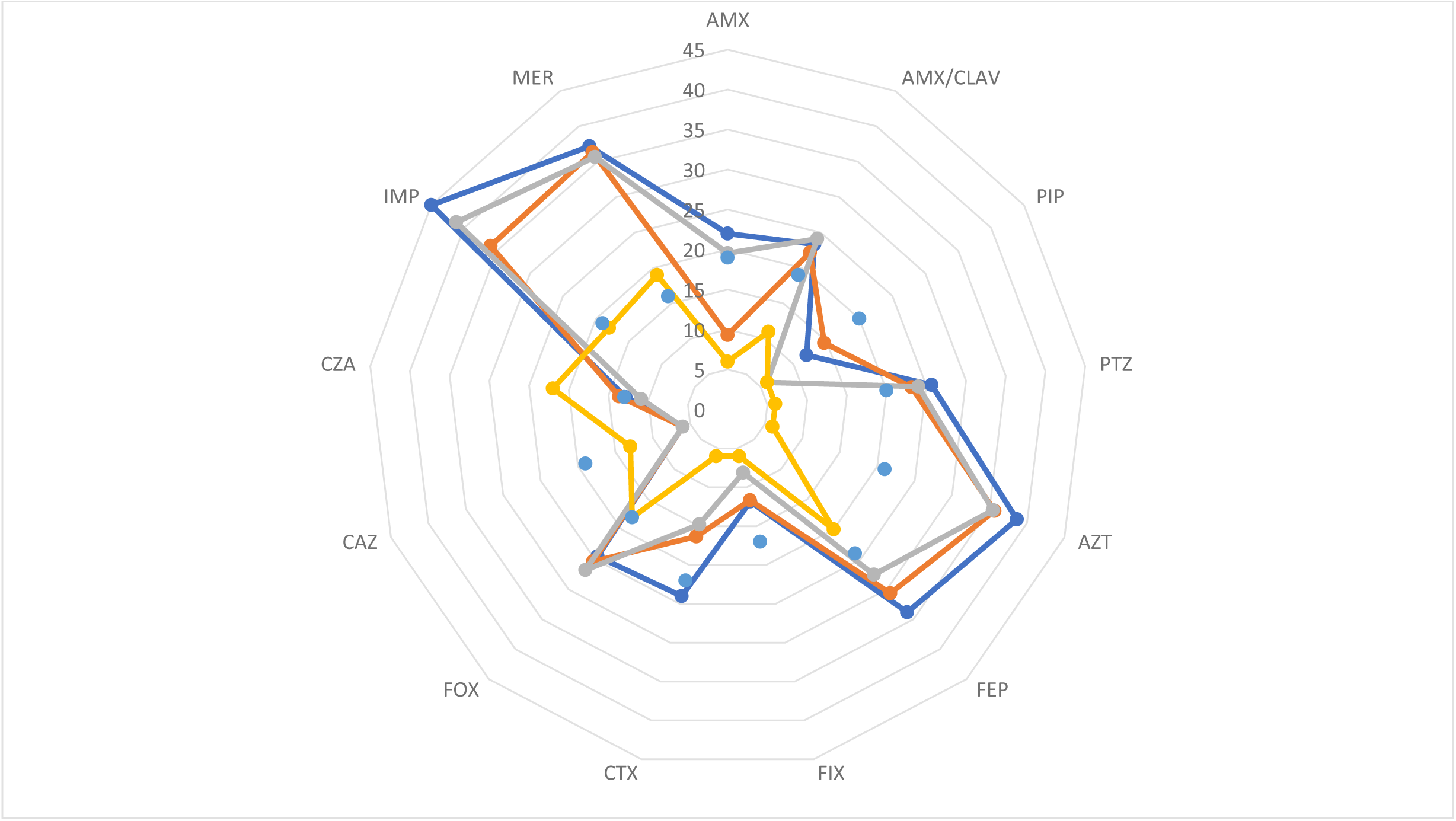
Kiviat diagram showing the distribution of mean inhibition zone diameters (mm) for the principal antibiotic molecules tested on selected variants (blue – variants enriched from KPC-2 (D179Y-D209V), orange – variants from KPC-3 (L169P), grey variants from KPC-33 (R6H, F20L +/- G291S) as well as ancestral KPC-2 variant (yellow). AMX, Amoxicillin; AMX/CLAV, amoxicillin/clavulanate; PIP, piperacillin; PTZ, piperacillin/tazobactam; AZT, Aztreonam; FEP, cefepime; FIX, cefixime; CTX, cefotaxime; FOX, cefoxitin; CAZ, ceftazidime; CZA, ceftazidime-avibactam; IMP, imipenem; MER, meropenem Diameters: millimeters; Blue points: EUCAST Breakpoints

Genotypic characterization: Sanger sequencing was performed on the 10 isolated colonies from each enrichment experiment. From KPC-2 mutants’ library, we identified only the combination of D179Y and D209V mutations on all tested mutants (n=10) which had cefiderocol MIC > 32 mg/L (corresponding to a >128-fold increase compared to wild-type KPC-2). From KPC-3 (H274Y), we identified only the L169P mutation (n=10) with cefiderocol MIC of 2 mg/L (17-fold increase compare to wild-type KPC-3) and from KPC-33 (D179Y) mutants’ library, we identified either the F20L (n=6), the F20L-G291S (n=2) or the R6H-F20L (n=2) mutations, all conferring a modest increase in cefiderocol MIC of 8 mg/L corresponding to a 4-fold increase compare to KPC-33.

To assess the direct impact of these mutations in the KPC gene on cefiderocol and other beta-lactams resistance, mutated pBR322 plasmids were extracted from selected strains. Antibiotic susceptibility testing and cefiderocol MIC were then determined after re- electroporating the plasmids into fresh *E. coli* TOP10 (Table 1). A decrease in cefiderocol MICs was observed following re-electroporation: from > 32 mg/L to 2 mg/L for KPC-2 with D179Y and D209V mutations, from 2 to 0.5 mg/L for KPC-3 with L169P and from 8 to 2 mg/L for KPC-33 (either with F20L, F20L-G291S or R6H-F20M mutations) while the susceptibility to other beta-lactams was not modified. This show that the KPC mutations alone do not fully explain the increase in cefiderocol MIC suggesting the contribution of other molecular mechanisms within the genomes to confer high levels of resistance to cefiderocol.

### Exploration of associated molecular mechanisms impacting Cefiderocol resistance

To explore the presence of underlying molecular mechanisms, WGS was performed on one representative isolate of each mutant libraries (KPC-2n2 from KPC-2, KPC-3n4 from KPC-3 and KPC-33n6 from KPC-33).

First, a snip comparison to the reference genome was performed. No newly acquired mutations were detected and notably in genes encoding porins, iron TonB-dependent transporters (TBDT) such as FepA, Fiu or CirA or penicillin-binding proteins (PBPs). In comparison to the reference genome, all analyzed genomes exhibited at least one insertion in the c*irA* gene, resulting in a truncated protein. In details, insertion sequences (IS) identified with ISFinder as ISKpn72, ISKpn8 and IS1X2 were present in *cir*A gene of KPC- 2n2, KPC-3n4 and KPC-33n6 isolates respectively. The integration occurred in the same region of the gene but at different sites.

In the KPC-2n2 we observed the insertion of another IS, ISKpn8 in the intergenic region between the *fiu* and *ybiX* genes (upstream of *ybiX)*, both being part of an operon.

### Deciphering the impact of each gene in cefiderocol MICs

To assess the contribution of *cirA, ybiX* and the combination of *cirA/ybiX* in cefiderocol resistance, MICs were assessed on Keio (*E. coli* K12) WT, and the same strain with Δ*cirA*, Δ*ybiX,* or *ΔcirA/ΔybiX* knockout mutants (the latter has been constructed for this study) with and without the pBR322-KPC-2n2 plasmid containing *bla*_KPC-2_ (D179Y-D209V), and pBR322-KPC-33 (D179Y) (Table 2).

**Table 2:** Cefiderocol MIC in *E. coli* keio wild-type strains as well as in Δ*cirA,* Δ*ybiX* and Δ*cirA*/Δ*ybiX strains with and without the bla*_KPC-2_ (with D179Y- D209V) or *bla*_KPC-33_ (D179Y) plasmids.

| Strains | Cefiderocol MIC (mg/L) | fold-increase in cefiderocol MIC compared to strains without plasmid | fold-increase in cefiderocol MIC compared to keio WT |
| --- | --- | --- | --- |
| keio WT | 0.03 | - | - |
| keio WT + *bla*<sub>KPC-2</sub> (with D179Y-D209V) | 0.5 | 16 | 16 |
| keio WT + *bla*<sub>KPC-33</sub> (D179Y) | 0.5 | 16 | 16 |
| keio Δ*cirA* | 0.12 | - | 4 |
| keio Δ*cirA* + *bla*<sub>KPC-2</sub> (with D179Y-D209V) | 2 | 16 | 66 |
| keio Δ*cirA* + *bla*<sub>KPC-33</sub> (D179Y) | 4 | 32 | 133 |
| keio Δ*ybiX* | 0.03 | - | - |
| keio Δ*ybiX* + *bla*<sub>KPC-2</sub> (with D179Y-D209V) | 1 | 32 | 32 |
| keio Δ*ybiX* + *bla*<sub>KPC-33</sub> (D179Y) | 0.5 | 16 | 16 |
| keio Δ*cirA*/Δ*ybiX* | 0.5 | - | 16 |
| keio Δ*cirA*/Δ*ybiX* + *bla*<sub>KPC-2</sub> (with D179Y-D209V) | >32 | 64 | 1066 |
| keio Δ*cirA*/Δ*ybiX* + *bla*<sub>KPC-33</sub> (D179Y) | >32 | 64 | 1066 |

Cefiderocol MICs for both Keio WT and Δ*ybiX* cells were initially very low (< 0.03 mg.L^-1^). In contrast the Δ*cirA* resulted in a higher baseline MIC (0.12 mg/L) compared to Keio WT. Upon introducing the *bla*_KPC-33_ (D179Y) or the *bla*_KPC-2_ (with D179Y-D209V) plasmids, the MIC increased from < 0.03 to 0.5 mg/L (for both plasmids) in Keio WT cells, from < 0.03 to 1 or 0.5 mg/L in Keio Δ*ybiX* cells and from 0.12 to 2 or 4 mg/L in Keio Δ*cirA* cells respectively for *bla*_KPC-33_ (D179Y) and the *bla*_KPC-2_ (with D179Y-D209V). The double deletion of *cir*A and *ybi*X led to a major increase in cefiderocol MICs from 0.5 mg/L without plasmid to > 32mg/L with both *bla*_KPC-33_ (D179Y) or the *bla*_KPC-2_ (with D179Y-D209V) plasmids (Table 2).

## Discussion

Using random mutagenesis, we introduced diversity into three KPC alleles: KPC-2, KPC-3, and KPC-33 (the latter being already described with increased cefiderocol MICs (18)) to evaluate mutational possibilities to confer high-level resistance to cefiderocol. Our mutagenesis allowed the generation of approximately 10⁴ to 10⁵ mutants per allele (with mean 3.6-mutations/alleles), corresponding in total to about 3-4 × 10⁵ different mutations. The hypothesis driving this work is that the accumulation of mutations in *bla*_KPC_ could enhance the cefiderocol resistance conferred by the KPC enzyme. A scenario similar to what was observed with TEM β-lactamases in response to ceftazidime a few decades earlier (23). Enrichment experiment under increasing concentrations of cefiderocol resulted in MIC increases, reaching >32, 2, and 8 mg/L for libraries derived from KPC-2, KPC-3, and KPC-33, respectively. When analysing the phenotypes of mutants selected at the highest cefiderocol concentrations, we observed a phenotypic convergence. The restoration of susceptibility to carbapenems among all mutants suggests an evolutionary trade-off, where the acquisition of specific mutations conferring resistance to cefiderocol may simultaneously enhance sensitivity to carbapenems and influence susceptibility to most other beta-lactams. Overall, the mutants exhibited resistance to ceftazidime and ceftazidime-avibactam, along with increased cefiderocol MICs. The cross-resistance to CZA and cefiderocol has recently been documented in the literature among clinical KPC variants, sometimes associated with other resistance mechanisms or beta-lactamases (17, 24, 25). Although these variants can be selected during CZA treatment, our results raise the concerning possibility of rapid selection of CZA resistant-mutants also under cefiderocol treatment (21, 24). This likely stems from shared structural features between ceftazidime and cefiderocol and a similar hydrolytic mechanism by KPC variants for the two drugs (22).

Among mutants derived from KPC-2 or KPC-3, only a single clone was enriched and subsequently dominated the population, as evidenced by identical sequences in all analysed mutants (10/10 clones). Interestingly, the D179Y mutation, frequently observed in clinical KPC variants and known to increase cefiderocol resistance, was selected in KPC-2 alongside the D209V mutation. The catalytic efficiency of KPC-2 has been described to be approximately 20 times lower for Cefiderocol compared to Ceftazidime (22). Nevertheless, the acquisition of the D179Y substitution significantly enhances the hydrolytic capacity of KPC-2 towards Cefiderocol by increasing the enzyme’s affinity for this antibiotic substrate. The D209V mutation is not located directly on the essential loops involved in the conformation of the active site of the enzyme, and has not been previously reported. To investigate the role of the D209V substitution in conferring high levels of resistance to Cefiderocol, Keio WT cells were transformed with the pBR322-KPC-33 plasmid (harboring only the D179Y mutation) and the pBR322-CKPC-2n2 plasmid (containing both D179Y and D209V mutations) and their MICs were assessed and compared. The Cefiderocol MICs were similar for both strains, suggesting that the D209V mutation did not contribute much to the increased Cefiderocol MIC. This highlights the adaptive importance of D179Y in cefiderocol resistance, as already suggested with clinical variants (6, 18). Of note, mutant libraries derived from KPC-33 (D179Y) did not yield other mutations that further enhanced significantly the resistance capabilities of the KPC enzyme.

Finally, the L169P substitution selected from KPC-3 is associated with a moderate increase in Cefiderocol MICs. This mutation has been previously reported in five clinical KPC variants and is often associated with other mutations (KPC-35, KPC-46, KPC-48, KPC-138, and KPC- 155) exhibiting varying resistance phenotypes to beta-lactams (26). Strains producing KPC- 35, KPC-46, and KPC-48 have been reported as susceptible to carbapenems, and KPC-46 and KPC-48 exhibit an ESBL-like resistance profile including resistance to ceftazidime-avibactam (6, 27, 28).

To disentangle the respective contributions of KPC mutations and chromosomal mechanisms to cefiderocol resistance, we assessed their individual impacts. For that, the impact of newly acquired KPC mutations on Cefiderocol resistance was subsequently assessed by isolating the plasmid containing the mutated *bla*_KPC_ gene from enriched mutant and reintroducing it into a naïve *E. coli* TOP10 strain, devoid of prior evolutionary changes. We found that the resulting Cefiderocol MICs attributed to the mutated KPC proteins, were elevated compared to the ancestral variants, though the increases were considerably more moderate (maximum an 8-fold increase with the plasmid from the KPC-2n2 mutant) (Table 1). These findings strongly suggest that additional molecular mechanisms are likely at play, and that chromosomal mutations are essential for achieving high-level cefiderocol resistance in these KPC variants. However, we started from most prevalent KPC variants (KPC-2, KPC-3, and KPC-33) but we cannot rule out that our results could have been different starting from other variants.

A comprehensive analysis of the mutants via WGS did not reveal any newly mutational events involving PBPs or outer membrane proteins, such as porin channels. Recent literature aligns with this perspective, indicating that mutations affecting genes encoding porins or PBP3 transpeptidases alone are insufficient to confer resistance thresholds to cefiderocol in Enterobacterales (29, 30). But WGS analysis and results from transformation of knockout mutants with plasmids harboring KPC variants strongly suggest the involvement of the c*irA* gene in the acquisition of *in vitro* resistance in our mutants. Indeed, the introduction of the pBR322-KPC-2n2 plasmid into Keio WT cells raised the Cefiderocol MIC from 0.03 to 0.5 mg/L corresponding to a 16-fold increase, showing a direct impact of the D179Y and D209V mutations on Cefiderocol resistance, though the MIC remained below the EUCAST resistance threshold (2 mg/L) (31). The deletion of *cirA* gene in Keio strain resulted in a 4-fold increase in cefiderocol MIC (0.12 vs. < 0.03 mg/Lin WT), suggesting that the absence of CirA transport restricts Cefiderocol uptake into the bacterial periplasm, thus elevating MIC values as already shown (32). Furthermore, the integration of the pBR322-KPC-2n2 plasmid into Keio *cir*A-deficient cells led to an even greater increase in resistance, with MICs rising 16-fold relative to Δ*cirA* cells (from 0,12 to 2 mg/L) and more than 66-fold compared to WT (from < 0,03 to 2 mg/L). These results are consistent with existing data, as TBDT are the primary targets for *in vitro* and *in vivo* resistance against cefiderocol in Enterobacterales (29, 33). Recent studies have demonstrated the impact of loss of function of CirA in acquiring high levels of resistance, particularly in Enterobacterales producing New Delhi Metallo-beta-lactamase (NDM) (16, 30, 34, 35). Moreover, a Chinese multicentric study suggested that restoring normal CirA function can reverse resistance in NDM-producing *E. coli* (30). However, few studies reported the selection of mutations targeting the *cirA* gene that confer significant Cefiderocol resistance in KPC-producing *E. coli*. *In vitro* study from Nurjadi et al. also highlighted that production of NDM facilitated the propensity for acquisition of *cirA* mutations, whereas KPC-2 did not (35). Our results suggest that the production of mutated KPC-2, KPC-3, and KPC-33 by *E. coli* strains may act synergistically in combination with mutations affecting the *cirA* gene to reach high cefiderocol MIC.

In parallel, WGS revealed the insertion of ISKpn8 within the intergenic region between the *fiu* and *ybi*X genes (upstream of the *ybi*X gene) of KPC-2n2. While the *fiu* gene is well-characterized and encodes one of the TBDTs essential for the transport of siderophores into the bacterial periplasm, the *ybiX* gene is described here for the first time as associated with cefiderocol resistance. This gene encodes a putative iron uptake factor and operates within an operon alongside *fiu*, which codes for the outer membrane iron-catecholate transporter, both regulated by a common transcriptional regulator : the Ferric Uptake Regulator (Fur) (36–40). *ybi*X is homologous to *piu*C in *Pseudomonas aeruginosa*, a gene implicated in siderophore uptake (41) which has been linked to *in vitro* resistance against siderophore-based antibiotics (42, 43). A similar mechanism is likely responsible for the observed resistance in the *E. coli* mutant, supporting the hypothesis that *ybi*X is implicated in the iron acquisition pathway. Recent literature indicates that the *in vitro* resistance level to cefiderocol is influenced by the contribution of each key TBDT involved in antibiotic substrate transport, and that the combined loss of Fiu function alongside a deletion of the *cirA* gene in *E. coli* significantly increase the MIC (32). Here, the KPC-2n2 strain exhibited the highest cefiderocol MIC – >32 mg/L and is the one that combined an insertion in both *cir*A and upstream of *ybiX* (distancing the promoter and thereby impacting expression) genes alongside with the expression of pBR322-KPC-2 with the D179Y and D209V mutations. The double deletion likely causes a strong restriction in cefiderocol uptake, explaining the marked increase in MIC.

Conclusion: In this study, we explored how libraries of mutants of KPC-2, KPC-3, or KPC-33 could adapt over a ten-day cefiderocol treatment. The resulting enriched mutants displayed similar phenotypes, including cross-resistance to both ceftazidime-avibactam and cefiderocol but with increased susceptibility to carbapenems and most other beta-lactams. While KPC mutations alone do not lead to high-level cefiderocol resistance in our experiment, we could hypothesize that they facilitate bacterial survival under cefiderocol treatment, creating opportunities for alternative resistance mechanisms, such as chromosomal mutations, to emerge. This intricate interplay between *bla*_KPC_ mutations and siderophore transport deficiencies underscores the importance of integrated therapeutic and surveillance approaches to mitigate resistance emergence and maintain effective treatment options. These findings provide critical insights into the resistance mechanisms of cefiderocol in KPC-producing *E. coli*. They suggest that the evolutionary pathway to high-level cefiderocol resistance is not straightforward, relying on both enzymatic mutations and chromosomal adaptations.

## Materials and methods

### KPC variants and mutagenesis experiments

*bla*_KPC-2_, *bla*_KPC-3_ (H274Y mutation) and *bla*_KPC-33_ (D179Y mutation) cloned into pBR322 plasmids were sourced from our existing collection (44) and used as templates for mutagenesis experiments. To explore KPC overall evolvability and diversity through random point mutations across the *bla*_KPC_ gene, we employed a PCR protocol using error-prone DNA polymerase (Mutazyme II DNA polymerase, GeneMorph II Random Mutagenesis Kit, Agilent Technologies). The protocol was performed as specified by the manufacturer and optimized to achieve mutation frequency of around 5 nucleotides changes per KPC genes (with a size close to one kb). Primers KPC-PCRmuta-F (5’- TAACCCTGATAAATGCTTCAATAATATTGAAAAAGGAAGAGTATG-3’) and KPC-PCRmuta-R (5’- TAAATCAATCTAAAGTATATGAGTAAACTTGGTCTGACAGTTA-3’) were used to amplify the *bla*_KPC_ gene. The mutated genes were then cloned using Gibson assembly (NEBuilderⓇ HiFi DNA Assembly, New England Biolabs) at a 5:1 ratio into the pBR322 plasmid, which had been pre-amplified with pBR322-pr-Gibson-F (5’- CATACTCTTCCTTTTTCAATATTATTGAAGCATTTATCAGGGTTA-3’) and pBR322-pr-Gibson-R (5’-TAACTGTCAGACCAAGTTTACTCATATACTTTAGATTGATTTA-3’) primers and purified using the QIAquick Gel Extraction Kit (Qiagen). After assembly, mutated plasmids were transformed by electroporation and expressed in One Shot™ TOP10 Electrocomp™ *E. coli* (Invitrogen) strains. Selection was performed on LB-tetracycline medium to generate the mutant libraries.

### *In vitro* enrichment experiment

The enrichment experiment was designed to select for mutants with high-level resistance to cefiderocol. A 10^7^ CFU/mL initial inoculum of the libraries of mutants were exposed to progressively increasing cefiderocol concentrations (ranging from 0.03 and up to 32 mg/L) in an iron-depleted Mueller-Hinton (MH) medium in microdilution plates (with 200 µl total volume). Each well was subcultured daily with an inoculum target of 10⁶ CFU/mL, allowing for a gradual escalation of cefiderocol concentration up to 32 mg/L. At the end of the ten-day period, viable resistant isolates that grew at the highest cefiderocol concentration were plated on LB-tetracycline medium, from which 10 colonies were randomly selected from each enrichment experiment and isolated for phenotypic and genotypic characterization.

#### Phenotypic characterization

The beta-lactams antibiotic susceptibility profile of the mutants were assessed both by the disk diffusion method on MH-agar medium and by broth microdilution plates using sensititre plates (ThermoFisher Scientific) in accordance with the last European Committee on Antimicrobial Susceptibility Testing guidelines (EUCAST) (31). Additionally, the cefiderocol MIC was determined using the Bruker’s UMIC® kit and CZA MIC was determined using the E-test® method (BioMérieux, Marcy l’Etoile, France) on MH-agar medium.

### Genotypic characterization of selected mutants

Sanger sequencing: for the selected mutants, the *bla*_KPC_ gene carried by pBR322 plasmid was amplified using standard PCR protocol and submitted for Sanger sequencing (GenewizⓇ Europe, Azenta Life Sciences) to identify mutations in the *bla*_KPC_ gene.

Whole genome sequencing: On ancestral strains as well as on one selected strain per enrichment experiment, genomic DNA extraction was performed using the Genomic DNA|gDNA Isolation Kits (Qiagen) for whole-genome sequencing (WGS). Libraries were prepared using the Illumina DNA Flex kit and sequenced on a NextSeq platform, using a NextSeq 500/550 Mid Output Kit v2.5 (300 cycles) (Illumina).

Bioinformatic analysis: Raw reads were trimmed using Trim Galore v0.4.4_dev and Trimmomatic v0.38 with a Phred score ≥ 20 and a minimum length of 50 bp. The quality of the trimmed reads was verified using FastQC v 0.11.8 and MultiQC v1.7. First, variant detection was performed based on the K12_DH5_alpha reference genome (GCF_002899475.1 NCBI accession). Alignment was carried out using BWA v 0.7.17-r1188, and variants were called using FreeBayes v 1.3.1-16. A custom python script was used to merge and generate a matrix of presence and absence of single nucleotide polymorphisms (SNPs). The pangenome was computed by constructing genomes of the samples with SPAdes v3.15.4. The presence and absence matrix was generated by Panaroo v1.5.0 based on Prokka v1.14 annotations. Both the SNPs and gene matrices were analyzed using an in-house R script to compare variations between cefiderocol resistant evolved strains and ancestral ones. Some insertions escaped detection by FreeBayes, particularly those with minimal overlap. To detect them, PanISa was used to specifically identify such cases and an R script was used to visualize the results (45).

### Determination of the role of mutational events in Cefiderocol resistance

To differentiate between resistance conferred by the selected KPC variants and potential other chromosomal mechanisms, we extracted mutated pBR322-KPC plasmids from the evolved strains, re-electroporated them into fresh One Shot™ TOP10 Electrocomp™ *E. coli*, and reassessed the cefiderocol MICs and antibiotic susceptibility testing.

Thanks to genomic analysis, we identified genomic modifications in some genes of the TBDT family (*cir*A gene and *ybi*X). To assess the contribution of these genes in Cefiderocol resistance phenotypes, cefiderocol MICs were assessed on strains from the Keio collection as well as on newly constructed deleted mutant (46): Keio WT, Δ*cirA*, Δ*ybiX, and ΔcirA/ΔybiX* knockout mutants before and after transformation with pBR322 containing the mutated *bla*_KPC._ For the construction of double-gene deletion mutant, the DNA fragments Δ*ybiX::kanamycin cassette (kmfrt)* was amplified from *E. coli* BW25113Δ*ybiX*::Kmfrt Keio strain. After removing the kanamycin cassette from *E. coli* BW25113Δ*cirA::*Km*frt* by transforming with the recombinase plasmid pCP20, the PCR products of Δ*ybiX::kmfrt* were introduced on the native chromosomal location under the native promoter in BW25113Δ*cirA::frt* using the λ-red linear recombinase plasmid pKOBEG to generate Δ*cirA/*Δ*ybiX* construct (47, 48). The integrity of all cloned fragments and mutations was verified by PCR with specific primers and DNA sequencing (Supplementary table 1).

## Acknowledgement

This research received no specific grant from any funding agency in the public, commercial, or not-for-profit sectors

---

**Supplementary Table 1:** Description of keio strains used and constructed for this study

| *E. coli* strains and plasmids | Genotype or description | Reference |
| --- | --- | --- |
| **Strains** |  |  |
| BW25113 | *E. coli* Wild-type strain | 1 |
| $\Delta cirA$ | BW25113$\Delta cirA::Kanamycin$ cassette (*kmfrt*) in the Keio collection | 1 |
| $\Delta ybiX$ | BW25113$\Delta ybiX::kmfrt$ in the Keio collection | 1 |
| $\Delta cirA/\Delta ybiX$ | Double mutant of $\Delta cirA$ and $\Delta ybiX$. BW25113$\Delta cirA$, $\Delta ybiX::kmfrt$ | This study |
| **Plasmids** |  |  |
| pCP20 | Rep(Ts) Flp+ | 2 |
| pKOBEG | oriR101ts araC arabinose-inducible $\lambda red$ $\gamma\beta\alpha$ operon | 3 |

- (1) Baba T, Ara T, Hasegawa M, Takai Y, Okumura Y, Baba M, Datsenko KA, Tomita M, Wanner BL, Mori H. 2006. Construction of Escherichia coli K-12 in-frame, gene knockout mutants: the Keio collection. Molecular systems biology 2:2006.0008. - (2) Cherepanov PP, Wackernagel W. 1995. Gene disruption in Escherichia coli: TcR and KmR cassettes with the option of Flp-catalyzed excision of the antibiotic-resistance determinant. Gene 158:9–14 - (3) Chaverroche MK, Ghigo JM, d'Enfert C. 2000. A rapid method for efficient gene replacement in the filamentous fungus Aspergillus nidulans. Nucleic acids research 28:E97

**Supplementary Table 2:**
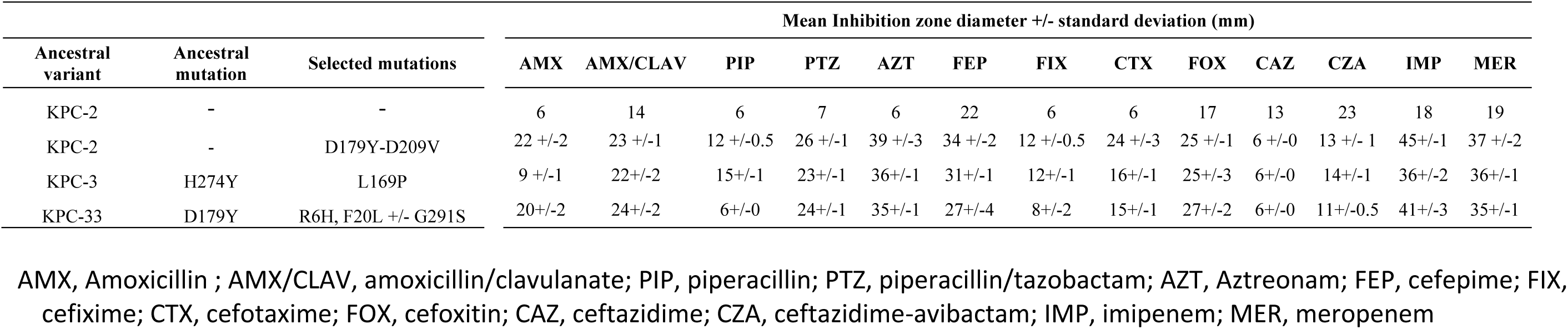
Results of antibiotic susceptibility tests performed by the disk diffusion method with representative beta-lactams (mean inhibition zone diameter +/- standard deviation in mm performed on 10 isolates)

## References

1. Logan LK, Weinstein RA. 2017. The Epidemiology of Carbapenem-Resistant Enterobacteriaceae: The Impact and Evolution of a Global Menace. J Infect Dis 215:S28–S36.

2. Cui X, Zhang H, Du H. 2019. Carbapenemases in Enterobacteriaceae: Detection and Antimicrobial Therapy. Front Microbiol 10:1823.

3. Karampatakis T, Tsergouli K, Lowrie K. 2023. Efficacy and safety of ceftazidime-avibactam compared to other antimicrobials for the treatment of infections caused by carbapenem-resistant *Klebsiella pneumoniae* strains, a systematic review and meta-analysis. Microb Pathog 179:106090.

4. Zhanel GG, Lawrence CK, Adam H, Schweizer F, Zelenitsky S, Zhanel M, Lagacé-Wiens PRS, Walkty A, Denisuik A, Golden A, Gin AS, Hoban DJ, Lynch JP, Karlowsky JA. 2018. Imipenem-Relebactam and Meropenem-Vaborbactam: Two Novel Carbapenem-β- Lactamase Inhibitor Combinations. Drugs 78:65–98.

5. Zhanel GG, Lawson CD, Adam H, Schweizer F, Zelenitsky S, Lagacé-Wiens PRS, Denisuik A, Rubinstein E, Gin AS, Hoban DJ, Lynch JP, Karlowsky JA. 2013. Ceftazidime-avibactam: a novel cephalosporin/β-lactamase inhibitor combination. Drugs 73:159– 177.

6. Hobson CA, Pierrat G, Tenaillon O, Bonacorsi S, Bercot B, Jaouen E, Jacquier H, Birgy A. 2022. Klebsiella pneumoniae Carbapenemase Variants Resistant to Ceftazidime-Avibactam: an Evolutionary Overview. Antimicrob Agents Chemother 66:e0044722.

7. Ding L, Shen S, Chen J, Tian Z, Shi Q, Han R, Guo Y, Hu F. 2023. *Klebsiella pneumoniae* carbapenemase variants: the new threat to global public health. Clin Microbiol Rev 36:e0000823.

8. Paul M, Carrara E, Retamar P, Tängdén T, Bitterman R, Bonomo RA, de Waele J, Daikos GL, Akova M, Harbarth S, Pulcini C, Garnacho-Montero J, Seme K, Tumbarello M, Lindemann PC, Gandra S, Yu Y, Bassetti M, Mouton JW, Tacconelli E, Rodríguez-Baño J. 2022. European Society of Clinical Microbiology and Infectious Diseases (ESCMID) guidelines for the treatment of infections caused by multidrug-resistant Gram-negative bacilli (endorsed by European society of intensive care medicine). Clin Microbiol Infect 28:521–547.

9. Tamma PD, Aitken SL, Bonomo RA, Mathers AJ, van Duin D, Clancy CJ. 2022. Infectious Diseases Society of America 2022 Guidance on the Treatment of Extended-Spectrum β- lactamase Producing Enterobacterales (ESBL-E), Carbapenem-Resistant Enterobacterales (CRE), and *Pseudomonas aeruginosa* with Difficult-to-Treat Resistance (DTR-P. aeruginosa). Clin Infect Dis 75:187–212.

10. Gaibani P, Amadesi S, Lazzarotto T, Ambretti S. 2022. Genome characterization of a Klebsiella pneumoniae co-producing OXA-181 and KPC-121 resistant to ceftazidime/avibactam, meropenem/vaborbactam, imipenem/relebactam and cefiderocol isolated from a critically ill patient. J Glob Antimicrob Resist 30:262–264.

11. Gato E, Guijarro-Sánchez P, Alonso-García I, Pedraza-Merino R, Conde A, Lence E, Rumbo-Feal S, Peña-Escolano A, Lasarte-Monterrubio C, Blanco-Martín T, Fernández- González A, Fernández-López MDC, Maceiras R, Martínez-Guitián M, Vázquez-Ucha JC, Martínez-Martínez L, González-Bello C, Arca-Suárez J, Beceiro A, Bou G. 2023. In vitro development of imipenem/relebactam resistance in KPC-producing *Klebsiella pneumoniae* involves multiple mutations including OmpK36 disruption and KPC modification. Int J Antimicrob Agents 62:106935.

12. Lombardo D, Ambretti S, Lazzarotto T, Gaibani P. 2022. In vitro activity of imipenem- relebactam against KPC-producing *Klebsiella pneumoniae* resistant to ceftazidime- avibactam and/or meropenem-vaborbactam. 5. Clin Microbiol Infect 28:749–751.

13. El-Lababidi RM, Rizk JG. 2020. Cefiderocol: A Siderophore Cephalosporin. 12. Ann Pharmacother 54:1215–1231.

14. Sato T, Yamawaki K. 2019. Cefiderocol: Discovery, Chemistry, and In Vivo Profiles of a Novel Siderophore Cephalosporin. Suppl 7. Clin Infect Dis 69:S538–S543.

15. Aoki T, Yoshizawa H, Yamawaki K, Yokoo K, Sato J, Hisakawa S, Hasegawa Y, Kusano H, Sano M, Sugimoto H, Nishitani Y, Sato T, Tsuji M, Nakamura R, Nishikawa T, Yamano Y. 2018. Cefiderocol (S-649266), A new siderophore cephalosporin exhibiting potent activities against *Pseudomonas aeruginosa* and other gram-negative pathogens including multi-drug resistant bacteria: Structure activity relationship. Eur J Med Chem 155:847–868.

16. Jousset AB, Poignon C, Yilmaz S, Bleibtreu A, Emeraud C, Girlich D, Naas T, Robert J, Bonnin RA, Dortet L. 2023. Rapid selection of a cefiderocol-resistant *Escherichia coli* producing NDM-5 associated with a single amino acid substitution in the CirA siderophore receptor. 4. Journal of Antimicrobial Chemotherapy 78:1125–1127.

17. Poirel L, Sadek M, Kusaksizoglu A, Nordmann P. 2022. Co-resistance to ceftazidime- avibactam and cefiderocol in clinical isolates producing KPC variants. Eur J Clin Microbiol Infect Dis 41:677–680.

18. Hobson CA, Cointe A, Jacquier H, Choudhury A, Magnan M, Courroux C, Tenaillon O, Bonacorsi S, Birgy A. 2021. Cross-resistance to cefiderocol and ceftazidime-avibactam in KPC β-lactamase mutants and the inoculum effect. Clin Microbiol Infect 27:1172.e7–1172.e10.

19. Castillo-Polo JA, Hernández-García M, Morosini MI, Pérez-Viso B, Soriano C, De Pablo R, Cantón R, Ruiz-Garbajosa P. 2023. Outbreak by KPC-62-producing ST307 Klebsiella pneumoniae isolates resistant to ceftazidime/avibactam and cefiderocol in a university hospital in Madrid, Spain. 5. J Antimicrob Chemother 78:1259–1264.

20. Amadesi S, Bianco G, Secci B, Fasciana T, Boattini M, Costa C, Gaibani P. 2024. Complete Genome Sequence of a *Klebsiella pneumoniae* Strain Carrying Novel Variant blaKPC-203, Cross-Resistant to Ceftazidime/Avibactam and Cefiderocol, but Susceptible to Carbapenems, Isolated in Italy, 2023. 6. Pathogens 13:507.

21. Giufrè M, Errico G, Del Grosso M, Pagnotta M, Palazzotti B, Ballardini M, Pantosti A, Meledandri M, Monaco M. 2024. Detection of KPC-216, a Novel KPC-3 Variant, in a Clinical Isolate of *Klebsiella pneumoniae* ST101 Co-Resistant to Ceftazidime-Avibactam and Cefiderocol. Antibiotics (Basel) 13:507.

22. Birgy A, Nnabuife C, Palzkill T. 2024. The mechanism of ceftazidime and cefiderocol hydrolysis by D179Y variants of KPC carbapenemases is similar and involves the formation of a long-lived covalent intermediate. Antimicrob Agents Chemother 68:e0110823.

23. Salverda MLM, De Visser JAGM, Barlow M. 2010. Natural evolution of TEM-1 β- lactamase: experimental reconstruction and clinical relevance. FEMS Microbiol Rev 34:1015–1036.

24. Fröhlich C, Sørum V, Tokuriki N, Johnsen PJ, Samuelsen Ø. 2022. Evolution of β- lactamase-mediated cefiderocol resistance. J Antimicrob Chemother 77:2429–2436.

25. Gaibani P, Ambretti S, Campoli C, Viale P, Re MC. 2020. Genomic characterization of a *Klebsiella pneumoniae* ST1519 resistant to ceftazidime/avibactam carrying a novel KPC variant (KPC-36). Int J Antimicrob Agents 55:105816.

26. Naas T, Oueslati S, Bonnin RA, Dabos ML, Zavala A, Dortet L, Retailleau P, Iorga BI. 2017. Beta-lactamase database (BLDB) - structure and function. J Enzyme Inhib Med Chem 32:917–919.

27. Hemarajata P, Humphries RM. 2019. Ceftazidime/avibactam resistance associated with L169P mutation in the omega loop of KPC-2. 5. J Antimicrob Chemother 74:1241–1243.

28. Cano Á, Guzmán-Puche J, García-Gutiérrez M, Castón JJ, Gracia-Ahufinger I, Pérez- Nadales E, Recio M, Natera AM, Marfil-Pérez E, Martínez-Martínez L, Torre-Cisneros J. 2020. Use of carbapenems in the combined treatment of emerging ceftazidime/avibactam-resistant and carbapenem-susceptible KPC-producing *Klebsiella pneumoniae* infections: Report of a case and review of the literature. J Glob Antimicrob Resist 22:9–12.

29. Kriz R, Spettel K, Pichler A, Schefberger K, Sanz-Codina M, Lötsch F, Harrison N, Willinger B, Zeitlinger M, Burgmann H, Lagler H. 2024. In vitro resistance development gives insights into molecular resistance mechanisms against cefiderocol. J Antibiot 10.1038/s41429-024-00762-y.

30. Wang Q, Jin L, Sun S, Yin Y, Wang R, Chen F, Wang X, Zhang Y, Hou J, Zhang Y, Zhang Z, Luo L, Guo Z, Li Z, Lin X, Bi L, Wang H. 2022. Occurrence of High Levels of Cefiderocol Resistance in Carbapenem-Resistant *Escherichia coli* before Its Approval in China: a Report from China CRE-Network. 3. Microbiol Spectr 10:e02670–21.

31. 2024. European Committee on Antimicrobial Susceptibility Testing. Data from the EUCAST MIC distribution website, last accessed 26 Sept 2024. https://www.eucast.org. Retrieved 26 September 2024.

32. Ito A, Sato T, Ota M, Takemura M, Nishikawa T, Toba S, Kohira N, Miyagawa S, Ishibashi N, Matsumoto S, Nakamura R, Tsuji M, Yamano Y. 2018. *In Vitro* Antibacterial Properties of Cefiderocol, a Novel Siderophore Cephalosporin, against Gram-Negative Bacteria. 1. Antimicrob Agents Chemother 62:e01454–17.

33. Klein S, Boutin S, Kocer K, Fiedler MO, Störzinger D, Weigand MA, Tan B, Richter D, Rupp C, Mieth M, Mehrabi A, Hackert T, Zimmermann S, Heeg K, Nurjadi D. 2022. Rapid Development of Cefiderocol Resistance in Carbapenem-resistant *Enterobacter cloacae* During Therapy Is Associated With Heterogeneous Mutations in the Catecholate Siderophore Receptor *cirA*. 5. Clinical Infectious Diseases 74:905–908.

34. Lan P, Lu Y, Chen Z, Wu X, Hua X, Jiang Y, Zhou J, Yu Y. 2022. Emergence of High-Level Cefiderocol Resistance in Carbapenem-Resistant *Klebsiella pneumoniae* from Bloodstream Infections in Patients with Hematologic Malignancies in China. 2. Microbiol Spectr 10:e00084–22.

35. Nurjadi D, Kocer K, Chanthalangsy Q, Klein S, Heeg K, Boutin S. 2022. New Delhi Metallo-Beta-Lactamase Facilitates the Emergence of Cefiderocol Resistance in *Enterobacter cloacae*. 2. Antimicrob Agents Chemother 66:e0201121.

36. Seo SW, Kim D, Latif H, O’Brien EJ, Szubin R, Palsson BO. 2014. Deciphering Fur transcriptional regulatory network highlights its complex role beyond iron metabolism in *Escherichia coli*. Nat Commun 5:4910.

37. Li Z, Pan Q, Xiao Y, Fang X, Shi R, Fu C, Danchin A, You C. 2019. Deciphering global gene expression and regulation strategy in *Escherichia coli* during carbon limitation. Microb Biotechnol 12:360–376.

38. Beauchene NA, Myers KS, Chung D, Park DM, Weisnicht AM, Keleş S, Kiley PJ. 2015. Impact of Anaerobiosis on Expression of the Iron-Responsive Fur and RyhB Regulons. 6. mBio 6:e01947–01915.

39. Panina EM, Mironov AA, Gelfand MS. 2001. Comparative analysis of FUR regulons in gamma-proteobacteria. Nucleic Acids Res 29:5195–5206.

40. Grinter R, Lithgow T. 2019. The structure of the bacterial iron-catecholate transporter Fiu suggests that it imports substrates via a two-step mechanism. J Biol Chem 294:19523–19534.

41. McHugh JP, Rodríguez-Quinoñes F, Abdul-Tehrani H, Svistunenko DA, Poole RK, Cooper CE, Andrews SC. 2003. Global iron-dependent gene regulation in *Escherichia coli.* A new mechanism for iron homeostasis. J Biol Chem 278:29478–29486.

42. Kim A, Kutschke A, Ehmann DE, Patey SA, Crandon JL, Gorseth E, Miller AA, McLaughlin RE, Blinn CM, Chen A, Nayar AS, Dangel B, Tsai AS, Rooney MT, Murphy-Benenato KE, Eakin AE, Nicolau DP. 2015. Pharmacodynamic Profiling of a Siderophore-Conjugated Monocarbam in *Pseudomonas aeruginosa*: Assessing the Risk for Resistance and Attenuated Efficacy. Antimicrob Agents Chemother 59:7743–7752.

43. McPherson CJ, Aschenbrenner LM, Lacey BM, Fahnoe KC, Lemmon MM, Finegan SM, Tadakamalla B, O’Donnell JP, Mueller JP, Tomaras AP. 2012. Clinically relevant Gram- negative resistance mechanisms have no effect on the efficacy of MC-1, a novel siderophore-conjugated monocarbam. Antimicrob Agents Chemother 56:6334–6342.

44. Hobson CA, Bonacorsi S, Hocquet D, Baruchel A, Fahd M, Storme T, Tang R, Doit C, Tenaillon O, Birgy A. 2020. Impact of anticancer chemotherapy on the extension of beta-lactamase spectrum: an example with KPC-type carbapenemase activity towards ceftazidime-avibactam. Sci Rep 10:589.

45. bvalot. 2024. bvalot/panISa. Python.

46. Baba T, Ara T, Hasegawa M, Takai Y, Okumura Y, Baba M, Datsenko KA, Tomita M, Wanner BL, Mori H. 2006. Construction of *Escherichia coli* K-12 in-frame, single-gene knockout mutants: the Keio collection. 1. Molecular Systems Biology 2:2006.0008.

47. Cherepanov PP, Wackernagel W. 1995. Gene disruption in Escherichia coli: TcR and KmR cassettes with the option of Flp-catalyzed excision of the antibiotic-resistance determinant. 1. Gene 158:9–14.

48. Chaveroche MK, Ghigo JM, d’Enfert C. 2000. A rapid method for efficient gene replacement in the filamentous fungus Aspergillus nidulans. 22. Nucleic Acids Res 28:E97.

